# Molecular prosthetics for CFTR designed for anion selectivity outperform amphotericin B in cultured cystic fibrosis airway epithelia

**DOI:** 10.1101/2025.08.28.671923

**Authors:** Jonnathan P. Marin-Toledo, Daniel Greenan, Nohemy Celis, Laura Haske, Agnieszka Lewandowska, Christopher K. Rakowski, Shashank Shastry, Arun Maji, Kelsie J. Green, Taras V. Pogorelov, Michael J. Welsh, Ian M. Thornell, Martin D. Burke

## Abstract

The ion channel-forming natural product amphotericin B (AmB) can serve as a molecular prosthetic for the cystic fibrosis transmembrane conductance regulator (CFTR) anion channel and thereby restore host defenses in cultured cystic fibrosis (CF) airway epithelia. This is despite the fact that the permeability of AmB-based channels favors cations, and these channels lose their capacity to increase airway surface liquid (ASL) pH in CF airway epithelia at high concentrations. We hypothesize that modifying such channels to favor anion permeability would make them more CFTR-like and thus increase their potential therapeutic effects compared to AmB. Here we show that a synthetic derivative of AmB, AmB-AA, which has an added positively charged appendage and forms ion channels with an improved relative permeability to anions, outperformed AmB in increasing the ASL pH in CF airway epithelia at both low and high concentrations. Further modifications led to another AmB derivative, C2’epiAmB-AA, that also minimized cholesterol binding and thus toxicity to cultured CF airway epithelia and was an effective surrogate for CFTR in primary cultured airway epithelia from people with CF.

## Introduction

Cystic fibrosis (CF) is a progressive disease caused by mutations in the CFTR gene leading to loss of CFTR anion channel function^1^. The ion channel-forming small molecule natural product, AmB, can serve as a molecular prosthetic for CFTR to partially restore HCO_3_^-^ and Cl^-^ secretion, airway surface liquid (ASL) pH, ASL hydration, and antibacterial properties in cultured CF airway epithelia, but only at low concentrations^2,3^. Unlike CFTR, AmB-based ion channels favor cation permeability^4,5^. We hypothesized that synthetically modifying AmB could yield derivatives that form channels with increased relative permeability for anions, making them more CFTR-like, and that such channels could have an improved therapeutic effect when compared to AmB^2,6^. Here we show that derivatives of AmB can be designed to have an improved relative permeability to anions, and that this can lead to more robust improvements in ASL pH in cultured CF airway epithelia which are maintained at both low and high concentrations.

The ion selectivity of biological protein channels is determined by factors such as pore diameter and electrostatic interactions, and the latter often highlighted as a primary contributor^7–9^. In the case of CFTR, there are two key aspects that influence anion selectivity. The cytosolic entry through the CFTR pore is lined with many positively charged lysine and arginine residues such as R128, K190, R303, K370, K1041, and R1048^9–11^. This creates an anion conducting pathway that stabilizes anions through electrostatic interactions. In addition, there is a narrow selectivity filter at the extracellular ends of transmembrane helices 1, 6, and 8 that contains residues G103, R334, F337, T338, and Y914 with the R334 residue being positively charged^10^. These two features collectively influence the anion selectivity of this protein channel [permeability of chloride relative to sodium (P_Cl_/P_Na_) ∼10-30]^12–14^.

Computational modeling predicts the C1-C13 polyol regions of multiple AmB molecules line an ion conducting pore^15–21^. These models also position the negatively charged C41 carboxylate near the channel entrance with the positively charged C3’ amine somewhat further away (**Fig. 1a**). A previous series of studies based on functional group deletions and/or other modifications of AmB provide support for this general channel architecture^15,16,22–25^. We also found that many AmB derivatives, other than those modified at C35,^15,25^ can still form channels^22–25^. We thus reasoned that converting the negatively charged C41-carboxylate group into a positively charged variant would yield a derivative that forms ion channels with improved relative permeability to anions^26–35^. Specifically, we added a positively charged ethylene diamine appendage to the C41 position of AmB (**Fig. 1a**), thereby mimicking the positive charges found lining CFTR and other protein anion channels.

**Figure 1.**
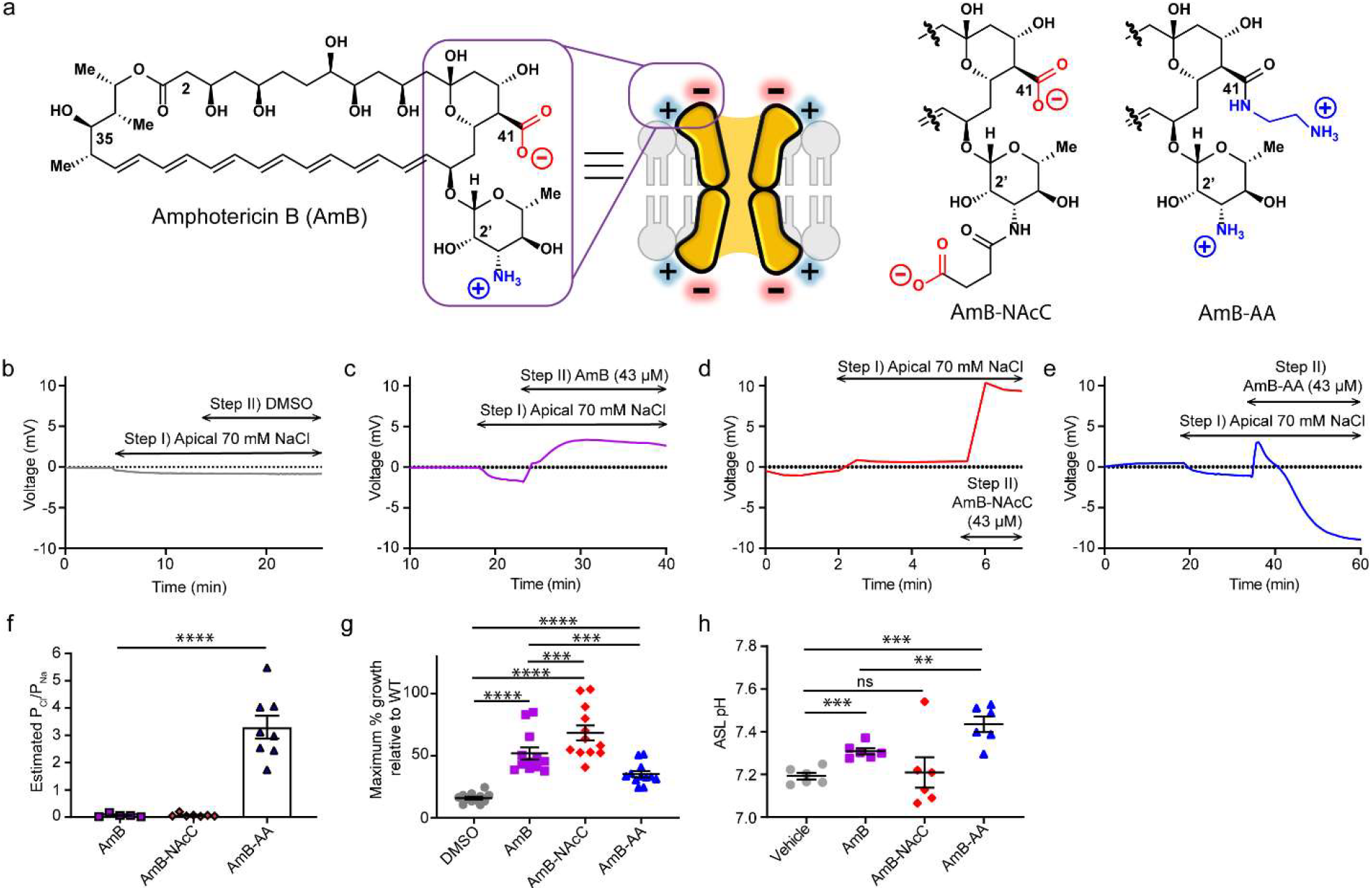
Differing ion selectivities for AmB-AA and AmB-NAcC channels are associated with different effects in preclinical models for protein channel deficiencies. (**a**) Chemical structure of AmB, C3’ N-Acetyl ethyl carboxylate (AmB-NAcC) and C16-Aminoethyleneamide AmB (AmB-AA). (**b**) Graphed representative traces of Ussing chamber current clamp ion-selectivity protocol prior to correcting for junction potentials. Step I) Apical media switch to 70 mM NaCl solution for paracellular ion movement determination. Step II) Addition of DMSO bilaterally. (**c**) Graphed results of Ussing chamber current clamp protocol done with 43 µM addition of AmB resulting in a positive change in voltage (cation selective). (**d**) Graphed results of Ussing chamber current clamp protocol done with 43 µM addition of AmB-NAcC resulting in a positive change in voltage (cation selective). (**e**) Graphed results of Ussing chamber current clamp protocol done with 43 µM addition of AmB-AA resulting in a negative voltage change (anion selective). (**f**) Estimated relative P_Cl_/P_Na_ for AmB, AmB-NAcC, and AmB-AA across FRT monolayers. (**g**) Maximum % growth relative to WT for all three compounds which was determined by comparing the trk1Δtrk2Δ and WT yeast growth across a concentration range for each compound. (**h**) ASL pH of CuFi-1 epithelial (dF508/dF508) treated with 2 µM AmB, AmB-NAcC or AmB-AA. (**f-g**) Data are represented as mean ± s.e.m. (**f**,**h**), One-Way ANOVA with multiple comparisons. (**g**) RM One-Way ANOVA with multiple comparisons. ns, not Significant. **** P <0.0001, *** P<0.001 ** P=0.0014, * P=0.01.

## Results

### AmB derivatives that favor anion flux

Towards our goal of identifying a molecular prosthetic that improves the relative permeability to anions, AmB was amidated with ethylenediamine to convert the negatively charged carboxylate at C41 into a positively charged amino ethyl appendage, yielding the bis-positively charged derivative, AmB-AA (**Fig. 1a**)^36–38^. As a control, a bis-negatively charged counterpart was also synthesized by acylating the C3’ amine with succinic anhydride yielding AmB-NAcC (**Fig. 1a**)^39^.

The relative permeability to Cl^-^ and Na^+^ (P_Cl_/P_Na_) for ion channels formed by AmB and these two synthetic derivatives were estimated by measuring dilution potentials (**Fig. 1b-1f**)^40,41^. Specifically, the transepithelial conductance (*G*_*t*_) value for an untreated Fischer rat thyroid (FRT) epithelium was obtained. Next, the apical [NaCl] was reduced to obtain a dilution potential for the epithelia. The ion channel-forming small molecule was then added to both sides of the epithelia to form conduits for ion flow and the experimental protocol was repeated to obtain *G*_*t*_ and a dilution potential for the monolayer treated with AmB or a derivative. With the *G*_*t*_ values and dilution potential values from this series of experiments, the *G*_*t*_ contribution by the small molecule was calculated and then the electrical relationship described by Equation 3 in the methods section was used to calculate the dilution potential produced by the small molecule. Finally, the P_Cl_/P_Na_ was calculated for each small molecule-based ion channel by substituting the small molecule dilution potential and known [NaCl] into the Goldman-Hodgkin-Katz equation (Equation 4 in methods section).

With this approach, it was estimated that AmB-based ion channels have a P_Cl_/P_Na_ of 0.06 ± 0.06 (**Fig. 1c, 1f**).^4,5^ The negative control, AmB-NAcC, showed a similar estimated P_Cl_/P_Na_ of 0.07 ± 0.06 (**Fig. 1d, 1f**). In contrast, the bis-positively charged derivative AmB-AA yielded a small molecule channel with a P_Cl_/P_Na_ of 3.0 ± 1.6 (**Fig. 1e, 1f**).

### Impacts of ion selectivity in channelopathy models

Having estimated that channels formed by AmB and AmB-NAcC favor cation permeability and those formed by AmB-AA have improved relative permeability to anions, we tested the effects of these three compounds in models of cation- and anion-channel deficiencies. In trk1Δtrk2Δ yeast that lack K^+^ transporting Trk1 and Trk2 proteins. AmB showed a maximum of 51.9% ± 4.85% growth relative to wild type (**Fig. 1g**)^42,43,44,45^. The AmB-NAcC ion channels were even more effective in restoring the maximum growth of trk1Δtrk2Δ yeast to 68.4% ± 6.03% of WT (**Fig. 1g and Extended Fig. 1**)^46–49^. In contrast, AmB-AA, which forms channels that have increased relative permeability to anions, was much less effective at recovering the growth of this cation channel-deficient yeast strain (maximum 35.2% ± 2.48% growth relative to WT) (**Fig. 1g**). AmB-AA also demonstrated a decreased capacity to increase intracellular K^+^ in trk1Δtrk2Δ yeast (**Extended Fig. 1**)^43,50^.

In cultured CF airway epithelial homozygous for the ΔF508 CFTR mutation (CuFi-1), AmB increased ASL pH at low concentrations (2 µM) (**Fig. 1h**)^2,3,51–55^. The bis-anionic derivative AmB-NAcC failed to increase ASL pH in this assay (**Fig. 1h and Extended Fig. 1**). However, the bis-cationic derivative, AmB-AA, caused a more substantial increase in ASL pH relative to AmB (**Fig. 1h**).

While AmB increases ASL pH in cultured CuFi-1 epithelia at low concentrations, this activity is lost at concentrations at or above 20 µM (**Fig. 2a**)^2^. In contrast, the anion-selective channel forming variant AmB-AA retained its capacity to increase ASL pH across the full range of tested concentrations (up to 100 µM) (**Fig. 2b**). We note that the net conductance for AmB- and AmB-AA-based channels at high concentrations is similar (**Fig. 2c**). These results collectively support the conclusion that AmB-AA-based channels, likely by increasing relative permeability to anions, outperform AmB-based channels as molecular prosthetics for CFTR in cultured CF airway epithelia.

**Figure 2.**
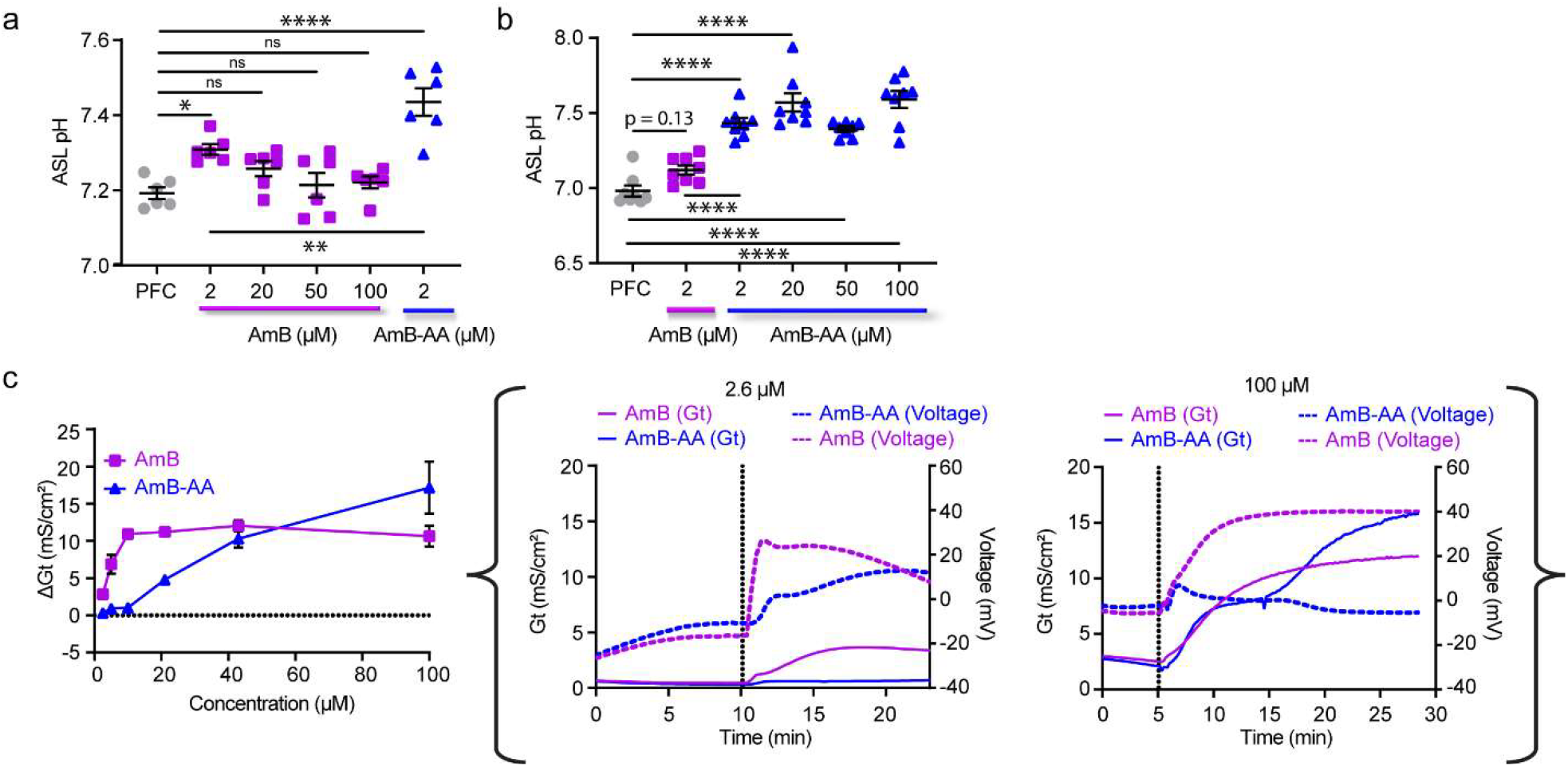
AmB-AA outperforms AmB as a Molecular Prosthetic. (**a**) ASL pH of CuFi-1 airway epithelial dose response between AmB at 0, 2, 20, 50,100 µM and AmB-AA at 2 µM (**b**) ASL pH of CuFi-1 airway epithelial dose response between AmB-AA at 0, 2, 20, 50, 100 µM and AmB at 2 µM (**c**) Conductance study with Fisher rat thyroid (FRT) mounted cells on Ussing chamber using apical 70 µM NaCl (pH=7) modified Ringer’s solution and basolateral 140 mM NaCl (pH=7) modified Ringer’s solution treated bilaterally with 2.69, 5.37, 10.75, 21.5, 43, or 100 µM of AmB, AmB-NAcC, and AmB-AA with complete Voltage (mV) and conductance (mS/cm^2^) traces for 2.6 µM and 100 µM runs. Data are represented as mean ± s.e.m. (**a**,**b**), Unpaired *t*-test. ns, not Significant. **** P <0.0001, *** P<0.001 ** P=0.0014, * P=0.01.

### Rationally minimizing toxicity

An orally inhaled dry powder formulation containing AmB, ABCI-003, was well-tolerated in clinical trials in healthy volunteers^56^, and in people with CF on and off modulators. These results support the use of AmB as a meaningful benchmark for minimum targeted tolerability in cultured airway epithelia. Although AmB-AA shows similar tolerability to AmB in a variety of mammalian cells (**Extended Fig. 2**), toxicity studies in cultured CuFi-1 and CuFi-4 airway epithelial treated with AmB and AmB-AA demonstrated increased toxicity for the latter (**Fig. 3a, 3b, and Extended Fig. 2d)**. We thus sought to mitigate this increased toxicity while keeping the anion selectivity of AmB-AA.

**Figure 3.**
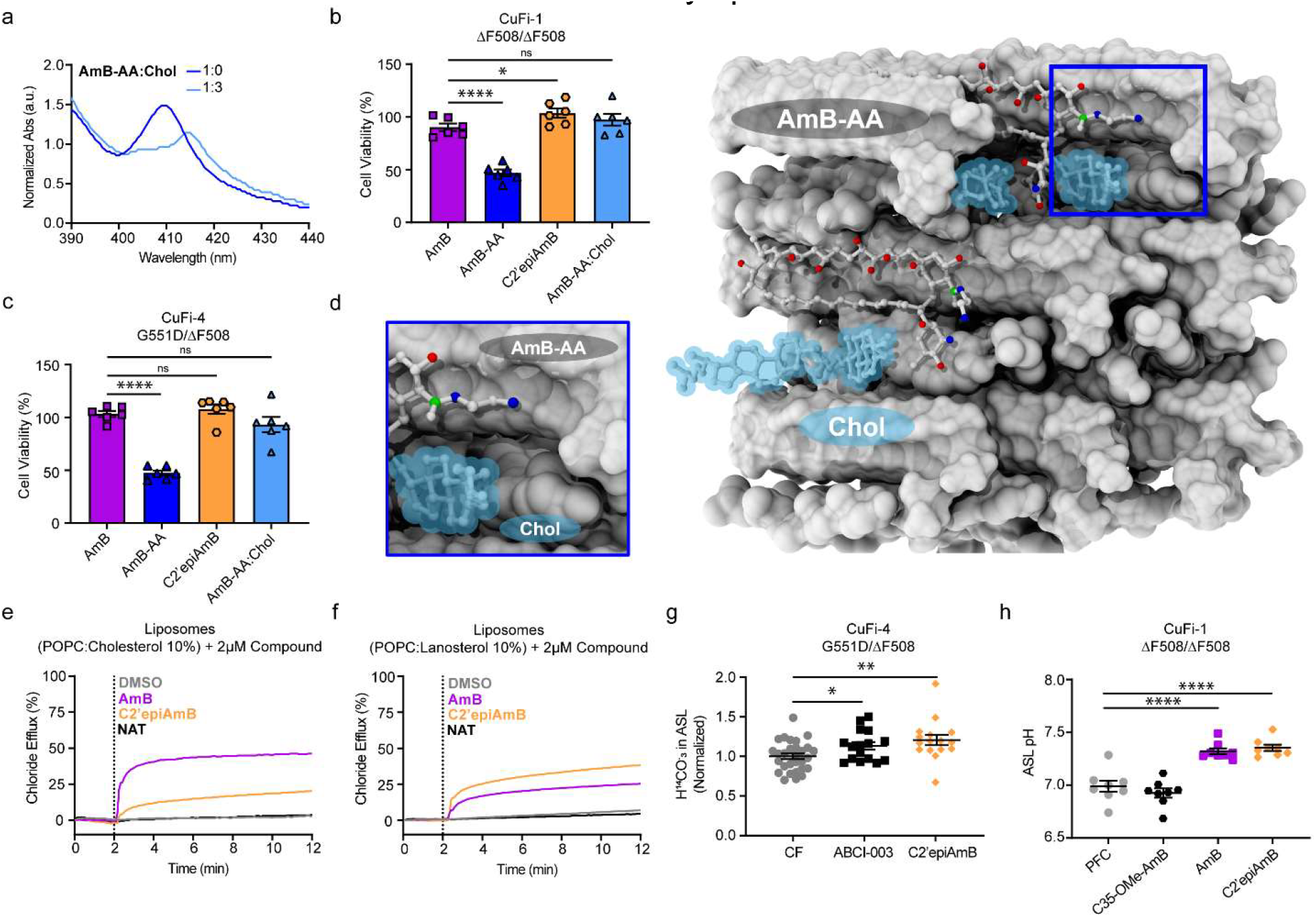
Toxicity of AmB-AA is mechanistically similar to AmB. (**a**) UV-Vis absorbance of AmB-AA with increasing ratios of cholesterol complexation. (**b**) Toxicity of AmB, AmB-AA, C2’epiAmB, and AmB-AA complexed with cholesterol all at 50 µM in CuFi-1 epithelial. (**c**) Toxicity of AmB, AmB-AA, C2’epiAmB, and AmB-AA complexed with cholesterol all at 50 µM in CuFi-4 epithelial. (**d**) Model of AmB-AA bound to cholesterol. (**e**) Chloride efflux from POPC liposomes with 10% cholesterol after DMSO, AmB, C2’epiAmB, and Natamycin addition. (**f**) Chloride efflux from POPC liposomes with 10% lanosterol after DMSO, AmB, C2’epiAmB, and Natamycin addition. (**g**) H^14^CO_3_ ^-^ transport of CuFi-4 epithelia after 48 hours of incubation with 50 µM ABCI- 003 and 50 µM C2’epiAmB. (**h**) ASL pH rescue of AmB, C2’epiAmB, and C35-OMe-AmB at 2 µM. Data are represented as mean ± s.e.m. ns, **(a**,**b**,**g**,**h)**, One-Way ANOVA with multiple comparisons. ns, not Significant. **** P <0.0001, *** P<0.001 ** P=0.0014, * P=0.01.

We previously demonstrated that the toxicity of AmB to human cells is linked to its ability to form an extramembraneous sterol sponge that binds and extracts cholesterol from mammalian lipid bilayers^25^. We first aimed to determine whether this sterol sponge mechanism was also primarily responsible for the toxicity of AmB-AA. Formation of an AmB-AA:Chol complex was confirmed by UV-Vis spectroscopy (**Fig. 3a**). Treating CuFi- and CuFi-4 airway epithelial with AmB-AA pre-complexed with cholesterol (AmB-AA:Chol) eliminated the toxicity to cultured airway epithelia (**Fig. 3b and 3c**). As expected, C2’epiAmB, a derivative known not to bind or extract cholesterol (**Extended Fig. 2f**)^24^, also caused no toxicity in cultured airway epithelia cell lines and showed reduced toxicity in other mammalian cell lines compared to AmB^25^ (**Fig. 3a, 3b, and Extended Fig. 2**).

Building on an extensive series of recently reported biophysical and structural studies with AmB^25,57^, XPLOR-NIH simulated annealing and molecular dynamics yielded a high-resolution structure of the AmB-AA sponge which showed the presence of a large void volume capable of accommodating the presence of a C16-ethylamine chain (**Fig. 4d**). These computational models further supported the conclusion that AmB-AA similarly kills mammalian cells via a sterol sponge-mediated cholesterol binding-dependent mechanism.

**Figure 4.**
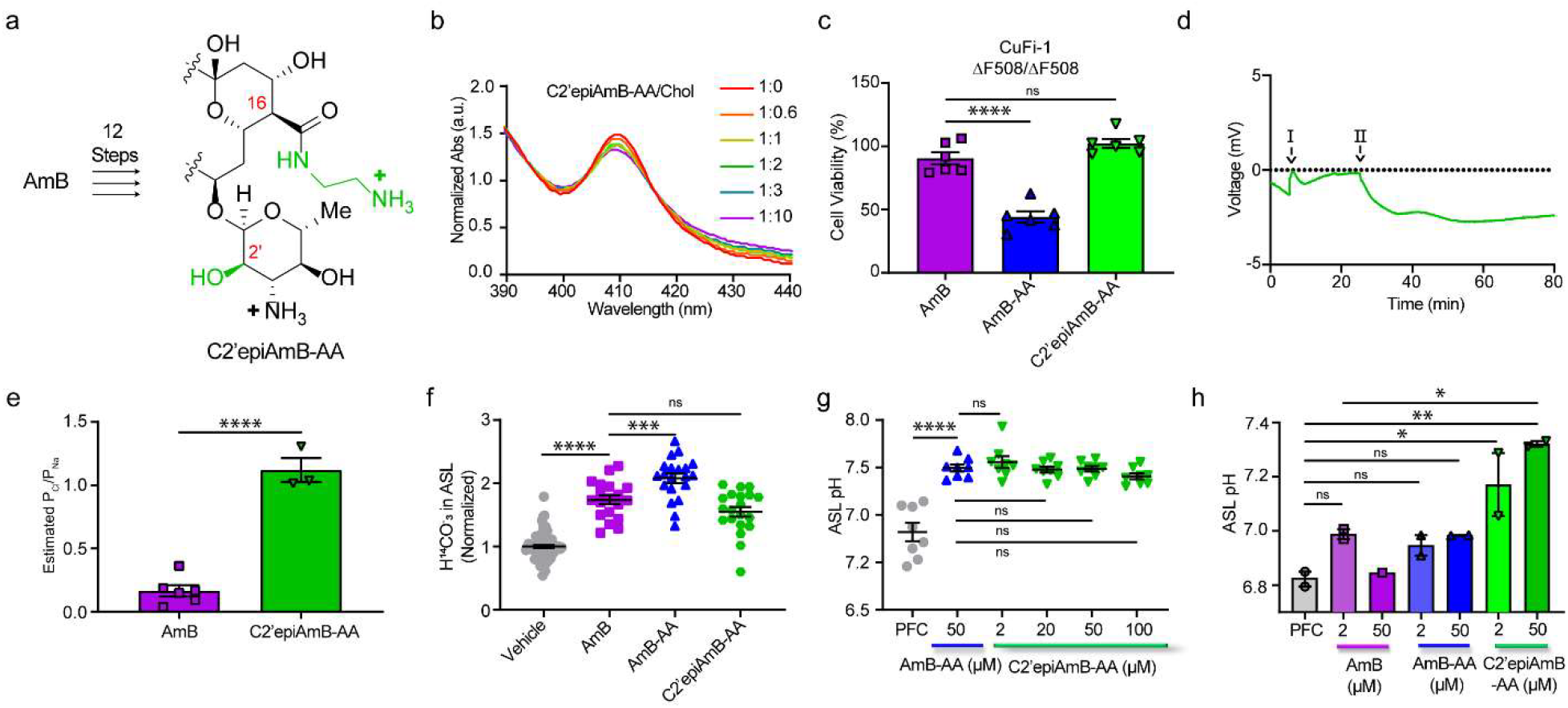
Epi-AmB-AA Retains Increased Anion Selectivity and is Less Toxic than AmB. **(a)** Synthetic scheme and chemical structure of C2’epiAmB-AA **(b)** UV-vis absorbance of C2’epiAmB- AA with increasing ratios of cholesterol **(c)** Toxicity of CuFi-1 cells treated with C2’epiAmB-AA, AmB, and AmB-AA at 50 µM **(d)** Graphed results of Ussing chamber current clamp ion-selectivity protocol. Step I) Apical media switch to 70 mM NaCl solution for paracellular ion movement determination. Step II) Addition of C2’epiAmB-AA AmB 100 µM bilateral **(e)** Estimated relative permeability of chloride to sodium ions for C2’epiAmB-AA at 200 µM and AmB at 44 µM across FRT monolayers **(f)** H^14^CO_3_ ^-^ rescue after 10 minutes of incubation with AmB, AmB-AA, and C2’epiAmB-AA at 2 µM in CuFi-4 airway epithelial **(g)** ASL pH dose response of C2’epiAmB-AA at 2, 20, 50, and 100 µM in CuFi-1 airway epithelial **(h)** ASL pH rescue or primary CF airway epithelial (both ΔF508/ΔF508) pretreated with 2 and 50 µM of AmB, AmB-AA, and Epi-AA. Data are represented as mean ± s.e.m. **(c-g)** One-Way ANOVA with multiple comparisons. **(h)** Unpaired *t*-test. ns, not Significant. **** P <0.0001, *** P<0.001 ** P=0.0014, * P=0.01.

Guided by our prior studies showing that epimerizing the C2’-OH of AmB mitigates cholesterol binding and thereby reduces the toxicity of AmB^25^, we tested whether we could reduce the toxicity of AmB-AA by making the same modification at C2’. Notably, while C2’epiAmB has been shown to form ion channels in the presence of ergosterol, it was not clear if C2’epiAmB or its derivatives that do not detectably bind cholesterol could form ion channels in cholesterol containing cells^25^.

Ussing chamber studies with FRT-derived monolayers failed to produce a G_t_ response with C2’epiAmB. However, in 10% cholesterol containing liposomes, 2 µM C2’epiAmB effluxed chloride as evidenced by an ion-selective electrode (**Fig. 3e**) although at a much lower level than AmB. Natamycin (NAT), a non-channel forming polyene macrolide natural product, showed no evidence of chloride efflux in this assay. This suggested that C2’epiAmB was forming ion channels in cholesterol-containing liposomes despite having no detectable binding to cholesterol. We next tested in sterol-free liposomes and found that both AmB and C2’epiAmB retained ion channel activity. The level of channel activity caused by AmB was significantly reduced relative to that seen in 10% cholesterol-containing liposomes. However, C2’epiAmB retained similar activity in the presence or absence of cholesterol (**Extended Fig. 3**). By repeating these efflux studies in 2, 5, 10, and 20% cholesterol-loaded liposomes, a clear cholesterol- dependent increase in permeabilizing activity for AmB was shown while C2’epiAmB was more or less unaffected by changes in membrane cholesterol content (**Extended Fig. 3**). These results suggested some capacity for these glycosylated polyene macrolide natural products to form ion channels in the absence of cholesterol binding.

We recognized, however, that sterol free liposomes can have different biophysical properties than their cholesterol-containing counterparts. Thus, we did the same experiment with 10% lanosterol containing liposomes which have been shown previously to have similar biophysical properties to ergosterol and cholesterol-containing liposomes^24,58–60^. Lanosterol does not detectably bind to AmB or C2’epiAmB, yet both of these polyene macrolides caused similar efflux as seen for each of them in sterol free liposomes (**Fig. 3f and Extended Fig. 3**). These results collectively support the conclusion that sterol binding is not required for ion channel formation by AmB and C2’epiAmB.

We next treated CuFi-4 airway epithelia with ABCI-003 and C2’epiAmB at 2 µM and quantified secretion of H^14^CO_3_^-^ (**Fig. 3g**). This confirmed that C2’epiAmB can transport H^14^ CO_3_^-^ to the ASL. We also treated cultured CuFi-1 airway epithelia with 2 µM C2’epiAmB, AmB, and a synthesized non-channel forming derivative of AmB, C35MeOAmB^15^. C35MeOAmB was not able to cause any detectable change in ASL pH. In contrast, C2’epiAmB successfully restored ASL pH to similar levels to that of AmB (**Fig. 3h**). These studies show that C2’epiAmB, which cannot detectably bind cholesterol, is able to form ion channels in cultured CF airway epithelia.

### C2’epiAmB-AA is a well-tolerated, more anion permeable MP for CFTR

The aforementioned studies suggested that epimerizing the C2’ position of AmB-AA might mitigate its toxicity while preserving its increased relative permeability to anions. Accordingly, amidation of the C2’epiAmB with ethylenediamine provided the bis-positively charged hybrid, C2’epiAmB-AA (**Fig. 4a**). As predicted, C2’epiAmB-AA showed no detectable cholesterol binding via UV-Vis spectroscopy (**Fig. 4b**) and, compared to AmB-AA and AmB, reduced toxicity in cholesterol containing cells (**Extended Fig.2**) including cultured CF airway epithelia (**Fig. 4c and Extended Fig. 2**). Unlike C2’epiAmB, C2’epiAmB-AA increased *G*_*t*_ in Ussing chamber studies. Dilution potential measurements in FRT monolayers revealed a higher P_Cl_/P_Na_ for C2’epiAmB-AA (1.12 ± 0.16) compared to that of AmB (0.16 ± 0.04). C2’epiAmB-AA increased bicarbonate secretion across CuFi-4 epithelia (**Fig. 4f**) resulting in an increase in ASL pH across all tested concentrations (**Fig. 4g**). Finally, primary airway epithelia cultured from people with CF were treated with 2 or 50 µM AmB, AmB-AA, or C2’epiAmB-AA. C2’epiAmB-AA caused a significant increase in ASL pH (**Fig. 4h**).

## Discussion

CFTR obtains its anion selectivity through positively charged amino acid residues that line the ion conduction pathways^9,10^. We added a positively charged ethylene diamine moiety near the putative small molecule channel entrance. This resulted in AmB-AA and C2’epiAmB-AA forming small molecule ion channels with increased relative permeability to anions.

There are three potential advantages for these new derivatives over AmB as molecular prosthetics to treat CF. First, AmB-AA increases ASL pH more effectively than AmB in cultured CF airway epithelia. Second, AmB-AA increases ASL pH even at high concentrations where AmB fails. Third, C2’epiAmB-AA should not extract native cholesterol from membranes and thus it should avoid potential disruptions in the activities of many transmembrane protein ion channels that depend on cholesterol for their functions^61–64^.

C2’epiAmB-AA thus represents a next-generation molecular prosthetic for CFTR with improved relative permeability to anions compared to AmB. This compound, or others with similar or further improved properties, may provide greater benefit to people with CF including those that cannot benefit from CFTR modulators. These findings may also have broader implications for designing optimized molecular prosthetics for other types of genetic diseases^65–67^.

## Methods

### General information

Unless mentioned otherwise, commercial reagents were stored and used as per the manufacturer’s recommendation. The sample size of the experiments was selected on the basis of literature precedence and preliminary experiments to adequately show the difference in outcome between different groups. No statistical methods were used to predetermine sample size. All experiments involving mammalian cell lines were carried out following a BSL2 safety protocol and the set up was approved by the Division of Research Safety, University of Illinois at Urbana-Champaign with primary lung epithelia experiments being performed at the University of Iowa. Fresh stock solutions of Amphotericin B and Amphotericin B derivatives were prepared in DMSO, and concentrations were determined by UV-vis. UV-vis specifications in methanol for AmB compounds include λ_max_ = 406nm, ε = 164,000 M^-1^ cm^-1^. Data and statistics were analyzed using Microsoft Excel for Windows (v.16.76 (23081101)) and GraphPad Prism 10 (v.10.5.0) on the Windows operating system. Chemical structures were drawn using Chemdraw 25.0.2. NMR data were analyzed using MestReNova 14.2.1. Final figures and images were processed using Adobe Illustrator 27.6.1.

### Materials

Commercially available materials were purchased from Sigma-Aldrich Co., GoldBio, Fisher Scientific, and Enamine and used without further purifications unless noted otherwise. AmB was purchased from Obiter Research LLC. Triethylamine was freshly distilled under nitrogen from CaH_2_. Water was double distilled or obtained from a Millipore MilliQ water purification system.

### Cell lines and Growth conditions

CuFi-1 and FRT cells (Welsh Laboratory, University of Iowa) were grown from cryostock on cell culture Treated 75 cm^2^ flasks (Thermo Scientific BioLite). The flasks for CuFi-1 cells were previously coated with 4 mL 50 μg/mL human placental collagen type IV (Sigma-Aldrich) for a minimum of 1 hours at 37 °C, then rinsed twice with PBS, and dried prior to seeding. Cells were cultured with 12 mL of Bronchial Epithelial Cell growth Medium (BEGM) with BulletKit (Lonza CC-3170) including eight SingleQuots of supplements (bovine pituitary extract (BPE), 2 mL; hydrocortisone, 0.5 mL; hEGF, 0.5 mL; adrenaline, 0.5 mL; transferrin, 0.5 mL; insulin, 0.5 mL; retinoic acid, 0.5 mL; triiodothyronine, 0.5 mL). The gentamycin-AmB supplement was discarded and was instead supplemented with 50 μg/mL penicillin–streptomycin (Corning Cellgro), 50 μg/mL gentamycin (Sigma-Aldrich G1397), and 2 μg/mL fluconazole (Sigma-Aldrich). Flasks for FRT cells were used without collagen pre-treatment. FRT cells were cultured with 12 mL of Ham’s F-12, 1X (Modified) with L-glutamine media (Corning 10-080-CV) supplemented with 50 μg/mL penicillin–streptomycin (Corning Cellgro), 50 μg/mL gentamycin (Sigma-Aldrich G1397), and 2 μg/mL fluconazole (Sigma-Aldrich). Both cell lines were grown to 90% confluence at 37 °C in 5% CO_2_, changing the medium every 2- 3 days, and then trypsinized with 4 mL 0.25% trypsin containing 1 mM EDTA (Gibco 25200-056). Trypsin was inactivated with 10 mL HEPES-buffered saline solution (Lonza CC-5024) with 1% fetal bovine serum (Gibco). Cells were spun down in an Eppendorf Centrifuge 5430R at 1,500 r.p.m. for 5 mins and resuspended in BEGM medium for passaging or Ultroser G medium for seeding. Seeding was done through resuspension of cells in 1:1 DMEM:Ham’s F-12 supplemented with 2% w/v Ultroser G (Crescent Chemical), 50 µg/mL penicillin-streptomycin (Corning Cellgro), 50 µg/mL gentamycin (Sigma-Aldrich G1397), and 2 µg/mL fluconazole (Sigma-Aldrich). CuFi-1 cell lines were seeded at 1.5 x 10^4^ cell per cm^2^ on 24 well Falcon plates with companion membranes 0.4-μm transparent PET membranes (Corning 3470). Membranes were coated with 100 µL collagen and then rinsed and dried similar to the flask protocol above. The cells were matured at an air-liquid interface by aspirating the apical surface 24 hours post-seeding and continuing basolateral media changes twice weekly for 14 days to reach full differentiation. Upon maturation, the cells were never used beyond 1.5 months (8 weeks post-seeding). FRT cell lines were seeded at 1.1 x 10^5^ cell per cm^2^ in non-collagen treated membranes in 24 well Falcon plates with companion membranes 0.4-μm Polycarbonate membranes (Corning 3413). The cells matured for 7 days while maintaining 150 µL of Ultroserg G medium on the apical membrane fresh. Upon maturation, the cells were never used beyond 4 days.

### Yeast Growth Conditions

Isogenic *Saccharomyces cerevisiae* BY4741(WT) and BYT12 (trk1Δtrk2Δ) were maintained on 15 mM KCl YPAD agar plates at 4 °C. Yeast was streaked from glycerol stocks and kept for 2-4 weeks. Overnight media was 100 mM KCl Yeast Nitrogen Base (YNB) pH 5.8 prepared from 4 g of (NH_4_)_2_SO_4_, 1.63 g of translucent K+ free YNB (ForMedium, CYN7505), 1.285 g of BSM (ForMedium, DBSM225), and 7.45 g KCl per liter. Media pH was adjusted to 5.8 with citric acid and ammonium hydroxide before autoclaving. Subsequently, 50 mL of sterile 40% w/v dextrose was added per liter (dextrose solutions were filter-sterilized using a 0.22 µm filter). The same formulation was followed for making K^+^-limiting media (15 mM KCl final) with 1.11 g of KCl added per liter.

### Small-Molecule Rescue Assay of Potassium-Transporter-Deficient Yeast

All experiments were done under sterile conditions. Overnight cultures of three to five cultures were grown to saturation over 12-24h, 200X RPM, 30 °C in 100 mM KCl YNB pH 5.8 media. Afterwards, 50 mL of cell suspension was pelleted at 1000XG, 23 °C for 5 min. Supernatant was removed, and cells were resuspended in 50mL of sterile Milli-Q H_2_O. Cells were centrifuged and washed 2 more times. After removing the supernatant, cells were resuspended in 15 mM KCl YNB, pH 5.8 and diluted with 15 mM KCl YNB media to OD_600_ = 0.090-0.110. Fresh stock solutions of Amphotericin B and Amphotericin B derivatives were prepared in DMSO, and concentrations were determined by UV-vis. UV-vis specifications in methanol for AmB compounds include λ_max_ = 406nm, ε = 164,000 M^-1^ cm^-1^. A 40-fold concentrated small molecule solution was prepared in DMSO for each treatment dose, and the small-molecule solution was delivered as a 1:40 v/v ratio for small molecule to cell suspension. Per tested concentration, 100 mL of cell suspension was treated with either a DMSO control or a small molecule in sterilized 250 mL flasks. Cells were grown protected from light for 12h at 200XRPM, 30 °C.

### Quantification of Total Cellular K^+^ Content via Inductively Coupled Plasma-Mass Spectroscopy

After 24h, 1 mL aliquots of each sample were utilized for OD_600_ values by UV-vis. Subsequently, samples were vacuum filtered through 0.8 µm membrane filters (EMD Millipore) and washed with 25 mL of ice-cold 20 mM MgCl_2_ (used ultrapure, molecular grade biology water). Yeast samples were resuspended in 5 mL of 20 mM MgCl_2_ and transferred into metal-free conical tubes, flash frozen, and lyophilized (≥72 hours). Next, under argon, dried samples were weighed and sealed in metal-free centrifuge tubes before inductively coupled mass spectroscopy.

### Ussing Chamber Dose-Response and Ion Selectivity Studies

Two NaCl solutions were prepared, one with a high [NaCl], and one with a low [NaCl]. The high [NaCl] solution contained 140 mM of NaCl, 5 mM HEPES, 1.2 mM calcium gluconate, 1.1 mM magnesium gluconate, and 5 mM glucose. The low [NaCl] Ringer’s solution contained was the same except for having a 70mM NaCl concentration. Solutions were titrated to a pH of 7.40 at 37 °C with NMDG and osmolarity was adjusted to 305-310 mOsm with mannitol using an Advanced Instruments Model 3250 Osmometer.

Mature FRT cells, prepared as mentioned above, were mounted in a Ussing chamber (Warner U2500) with 3M KCl agar bridges connected to amplifiers recording open-circuit trans epithelial voltage (V_t_) with intermittent current injections of ±0.5 nA. Membranes were mounted on the Ussing Chamber using the culture cup insert for Transwell adaptor, 6.5 mm (Warner U9924T-06). After completing dilution potential experiments on FRT monolayers, the monolayer was lysed with water and junction potentials induced by ion dilutions were assessed then subtracted from obtained dilution potentials.

For dose-response conductance studies (**Fig. 2d**): The cells were treated to 5 mL Apical low [NaCl] solution and 5 mL Basolateral High [NaCl] prior to starting conductance recording and allowed to equilibrate prior to the addition of DMSO, AmB, AmB-NAcC, or AmB-AA at selected concentrations of 2.69, 5.37, 10.75, 21.5, 43, or 100 µM. Conductance readings for each compound and concentration were summarized in **Fig. 2d** by determining the difference in conductance post and prior compound addition.

For ion selectivity studies (**Fig. 1b-1f and 4d**): The cells were treated to 5 mL bilateral High [NaCl] buffer. After equilibration, the apical solution was aspirated and replaced with 5 mL low [NaCl] buffer (**Fig. 1b-1f and 4d** step I). After equilibration of FRT paracellular ion flow, the cells were treated with either DMSO, AmB, AmB-NAcC, or AmB- AA at 43 µM and allowed to equilibrate (**Fig. 1b-1f and 4d** step II).

### Sodium and Chloride Activity Coefficients

Determining the activity coefficients (α_ion_) for sodium and chloride were calculated using the ionic strength of each solution and the extended Debye-Hückel equation^68^, which is applicable for physiological concentrations:

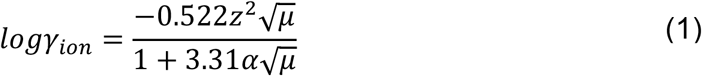

Where *γ* is the activity coefficient of the ion, *z* is the ionic charge, *µ* is the ionic strength of the solution, and α is the effective diameter of the hydrated ion^69^. The constants - 0.522 and 3.31 were used because experiments were performed at 37 °C^70^.

The ion activity was then calculated by the equation:

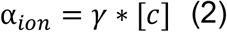

Where α is the ion activity, *γ* is the activity coefficient (α_Na_ =0.735872606 and α_Cl_ =0.71537313), and [c] is the ion concentration (140 mM).

### Permeability Calculations

We modeled the FRT monolayer as an electrical circuit so that:

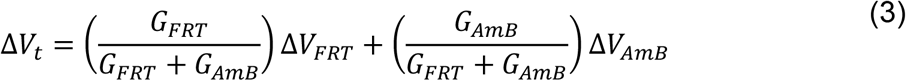

Where ΔV_t_ is the total measured dilution potential in the presence of small molecule, *G*_*FRT*_ is the trans epithelial conductance of the untreated FRT monolayer, G_FRT_ + G_AmB_ is the trans epithelial conductance of the FRT monolayer treated with an AmB-based small molecule, *V*_*FRT*_ is the measured dilution potential of the FRT monolayer prior to applying the AmB channel, G_AmB_ is obtained from subtracting the untreated monolayer *G*_*t*_ (*G*_*FRT*_) from the AmB-treated monolayer (G_FRT_ + G_AmB_), and *V*_*AmB*_ is the calculated the contribution of the small ion AmB pathway to *V*_*t*_.

*V*_*AmB*_ then can be used to calculate the relative permeability (P_Cl_/P_Na_) of the AmB small molecule using the Goldman-Hodgkin-Katz equation^71,72^:

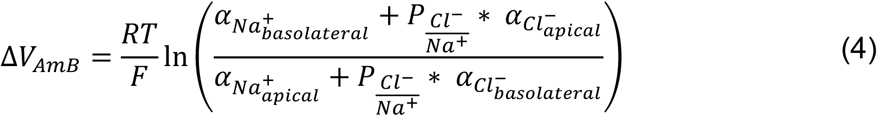

Approximating the constants at 37°C simplifies to:

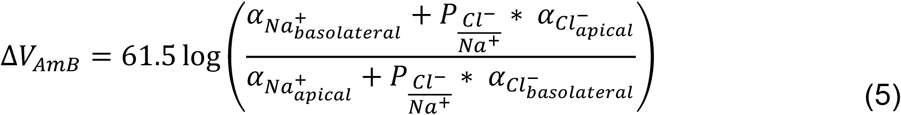

Where *F* is Faraday’s constant, *R* is the gas constant, *T* is temperature, *ΔV*_*AmB*_ is the dilution potential for the contribution of the small ion AmB pathway to *V*_*t*_, and P_Cl-/Na+_ is the relative permeability Cl^-^ to Na^+^.

### Data Analysis

Electrophysiological data was analyzed using a custom graphical user interface coded in MATLAB version R2018b (Mathworks). Equations were solved in MATLAB version R2018b (Mathworks) using custom code. All codes were written by Ian M. Thornell and are freely available upon request and the data that supported the findings of this study are available from the corresponding authors upon reasonable request.

### Airway Surface pH Studies

Determination of the ASL pH rescue was done following the same protocol previously reported^2^.

### H^14^ CO_3_^-^ Transport across CuFi monolayers

Small-diameter CuFi-4 cultured epithelia were used for this experiment (0.33 cm^2^). ^14^C-labelled sodium bicarbonate was obtained as a sterile 35.7 mM aqueous solution pH 9.5 (PerkinElmer). All experiments were run less than two months after seeding. Fresh USG medium was added to the basolateral side before experimentation. The apical membrane was treated with 20 μL vehicle, AmB (50 µM), or AmB derivative (50 µM )/ABCI-003 (50 µM), and the cultured epithelia were incubated for 48 h, at 37 °C in a 5% CO_2_ atmosphere. After the end of the treatment period, 5 μL of 1.4 mM H^14^ CO_3_^-^ stock solution in USG medium was added to the basolateral medium. The cultured epithelia were then incubated at 37 °C for 10 min. After 10 min, the apical membrane of the cultured epithelia was immediately washed with 200 μL PBS. 150 µL of the ASL wash was then collected and placed in 3 mL of Opti-Fluor (a high flash point LSC-cocktail for aqueous samples, Thermo Scientific). As a positive and negative control, 150 µL of the basolateral media containing ^14^C-labelled sodium bicarbonate or PBS was placed in Opti-Fluor. All samples were analyzed by liquid scintillation counting. The measured rate of basolateral-to-apical H^14^ CO_3_^-^ secretion was then normalized to the vehicle-treated control.

### CuFi-1/CuFi-4 alamarBlue Toxicity Studies

All experiments were run less than two months after seeding. Fresh USG medium was added to the basolateral side before experimentation. To cultured CuFi-1 or CuFi-4 epithelial 2.5 µL of a solution of desired compound in DPBS (400 µM stock; for a final concentration of 50 µM) was added to the apical membrane. The cells were then incubated for 48 h at 37 °C in a 5% CO_2_ atmosphere. After the end of the treatment period 2.5 µL of alamarBlue (ThermoFisher) was added to the apical membrane. Cells treated with 20 µL of Triton X were used as a control for 100% cytotoxicity. Using a plate reader, the emission at 585 nm was tracked using an excitation wavelength of 555 nm. The percentage viability was quantified by using the difference between compound viability, 100% viability (blank control), and 0% viability (Triton X-100 control).

### Primary human renal proximal tubule epithelial cell viability assay

Primary Renal cell viability assay was carried out as previously reported^25^.

### Human hepatocellular epithelial cell viability assay

Human hepatocellular epithelial cel viability assay was carried out as previously reported^25^.

### Sterol UV-Vis binding Assay

Sterol UV-Vis binding assay was carried out as previously reported^25^.

### Liposomes Preparation

A 7 mL vial was charged with 640 µL of POPC solution (25mg/mL of POPC in CHCl_3_) (For sterol free liposomes). For 10% cholesterol loaded liposomes a 230 µL solution of cholesterol (4mg/mL of recrystallized cholesterol in CHCl_3_) was added. For 10% lanosterol loaded liposomes a 230 µL solution of cholesterol was added instead. NOTE: This ratio was modified accordingly for the different percentages of sterol used for the dose response. To have enough liposomes to run two replicates, three 7 mL vials were prepared. The solvent was removed with gentle stream of nitrogen. The films were stored under high vacuum for 8+ hours prior to use. The films were then resuspended in 1 mL of a 250 mM NaCl, 40 mM HEPES buffer (pH 7.5) solution and vortexed for 3 minutes to get multilamellar vesicles (MLVs). The suspension was pulled into a Hamilton (Reno, NV) 1 mL gastight syringe. This syringe was put in a mini-extruder (Avanti Research, AL; Product # 610000) where the lipid solution was passed through a 0.20 µm Millipore (Billeric, MA) polycarbonate filter 21 times to isolate LUVs. These steps were repeated with every prepared vial. Each vial corresponding to one type of liposome were combined after extraction. The combined liposome solutions, sterol-free, cholesterol-loaded, or lanosterol-loaded, were then purified via dialysis using 3,500 dialysis cassettes. The solutions were dialyzed three times against 600 mL of a 62.5 mM MgSO_4_, 40 mM HEPES buffer solution (pH 7.3). The first two dialyses were 2 hours long meanwhile the last one was performed overnight.

### Liposomes Quantification

10 µL of the individual LUVs were added to 7 mL vials (in triplicate). 450 µL of H_2_SO_4_ (8.9 M) was added to each vial, including 3 additional 7 mL vials with no sample (blank control). The vials were then added to a 225 °C heated aluminum block for 25 minutes and then moved to room temperature and allowed to cool for 5 minutes. After cooling 150 µL of 30% w/v aqueous hydrogen peroxide was added to every vial and returned to the 225 °C heated aluminum block for 30 minutes. Samples were again moved to room temperature and allowed to cool for 5 minutes. Once cooled 3.9 mL of MilliQ water and 500 µL of 2.5% w/v ammonium molybdate solution was added along with 500 µL 10% w/v ascorbic acid. The vials were sealed with a PTFE-lined cap and vortex vigorously for 5 minutes before moving to a 100 °C heated aluminum block for 7 minutes. The vials were then moved to room temperature and allowed to cool for 15 minutes before 200 µL was taken, added into a 96-well plate and analyzed by UV/Vis spectroscopy (820 nm absorbance) using a plate reader. The concentration of liposomes was quantified by generating a standard curve by adding 0 (blank), 20, 40, 60, 80, 100, 120, 140 µL of a 0.65 mM phosphorus standard solution (Sigma) to 7 mL vials (in triplicate) and repeated all previous steps. Using this standard curve the concentration of phosphorous was quantified for all liposomes, for 3 vials per liposomes batch the concentration was around 13-17 mM.

### Chloride Efflux Experiments

Desired liposomes were diluted to 1 mM using a 62.5 mM MgSO4, 40 mM HEPES buffer solution (pH 7.3). 4 mL of this solution was added to a 20 mL vial (with 40 µL of ISA) and stirred. A chloride ion selective electrode (Fischer) was calibrated using standard Fischer protocols and then added ito the stirring mixture. Using the StarCom software, the baseline chloride ppm was recorded over a period of 2 minutes, followed by the addition of 40 µL of either DMSO or desired compound (0.2 mM compound stock solution in DMSO for a final concentration of 2 µM) and recorded the ppm change for a period of 10 minutes. To finish the readings, 40 µL of 30% v/v Triton X- 100 was added to the stirring mixture and the data was collected for an additional 3 minutes before stopping (total run time of 15 minutes). Percent chloride efflux was quantified by baselining the data using the first 2 minutes of the run and quantifying the change in ppm using the post Triton X-100 readings as 100% chloride efflux.

### System Preparation and Molecular Dynamics Simulations

All molecular simulations were performed using NAMD3^73^ and subsequent visualization done with VMD [ http://www.ks.uiuc.edu/Research/vmd/]^74^. The system was prepared using the lowest-energy AmB–Erg lattice (Fig. 1a) from previously reported data. The structure for the original AmB-Erg lattice4 was used as a template for modeling the AmB-AA + Sterol lattices. To maintain the structure of the experimentally resolved sponge, the macrolide region of all AmB molecules were constrained using the fixedAtoms flag in NAMD3 and all sterol molecules left unconstrained. AmB parameters were obtained from CHARMM General Force Field^75^ and sterols parameters were obtained from CHARMM36 forcefield^76^. After constructing each AmB-AA system, the sponge was solvated using the TIP3P water model^77^ and ionized at 150mM NaCl. An initial energy minimization was done for 10000 steps and a 1ns NPT equilibration run allowing water molecules to relax and system size to converge. For equilibration and production runs, the Nosé-Hoover constant pressure method^78,79^ was used to maintain constant pressure while temperature was maintained through langevin dynamics with a damping coefficient of 0.5 ps^−1^. The simulation was carried out using an isothermal–isobaric ensemble at 310 K with a pressure of 1.0 atm. A 12-Å cut-off was used for nonbonded interactions with a smoothing function implemented farther than 10 Å. The integration time step of 2 fs was used, and the bond distances of the hydrogen atoms were constrained using the SHAKE algorithm^80^ 29. For long-range electrostatic calculations, the particle mesh Ewald method30 was used, with a grid density of less than 1 Å−3. Production runs were 500 ns long with 3 replicates each.

## Acknowledgements

We gratefully acknowledge Rebecca Sponenburg for assistance with inductively coupled plasma-mass spectroscopy which was performed at the Northwestern University Quantitative Bio-element Imaging Center. We also acknowledge Corinne P Soutar for helpful discussions regarding CF assays. This work was supported by research funding from Cystetic Medicines (102351) and from the NIH (Grant R35 GM118185 to M.D.B; HL152960-05 to M.J.W). We also thank the NIH for funding support for N.C. a Predoctoral Fellow (5T32 GM136629-04).

## Extended Data

**Extended Figure 1:**
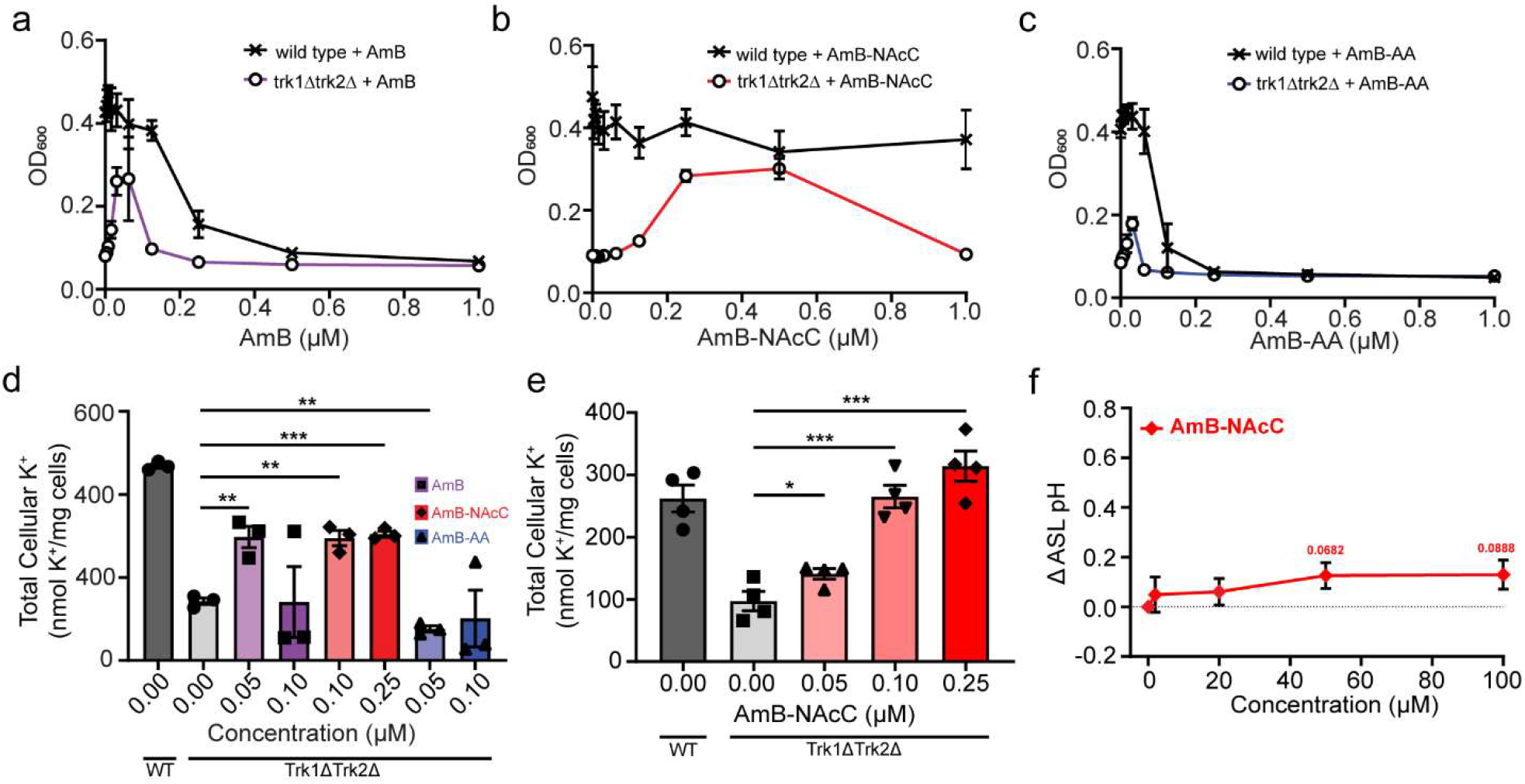
AmB-NAcC hints at higher cation selectivity than AmB in physiological Assays. **(a)** Dose response of AmB in yeast rescue assay. **(b)** Dose response of AmB-NAcC in yeast rescue assay. **(c)** Dose response of AmB-AA in yeast rescue assay. **(d)** Total Cellular K^+^ levels of yeast for AmB, AmB-AA, AmB-NAcC via ICPMS. **(e)** Total Cellular K^+^ levels of trk1Δtrk2Δ with AmB-NAcC at varying doses. **(f)** ASL pH dose response for AmB-NAcC.

**Extended Figure 2:**
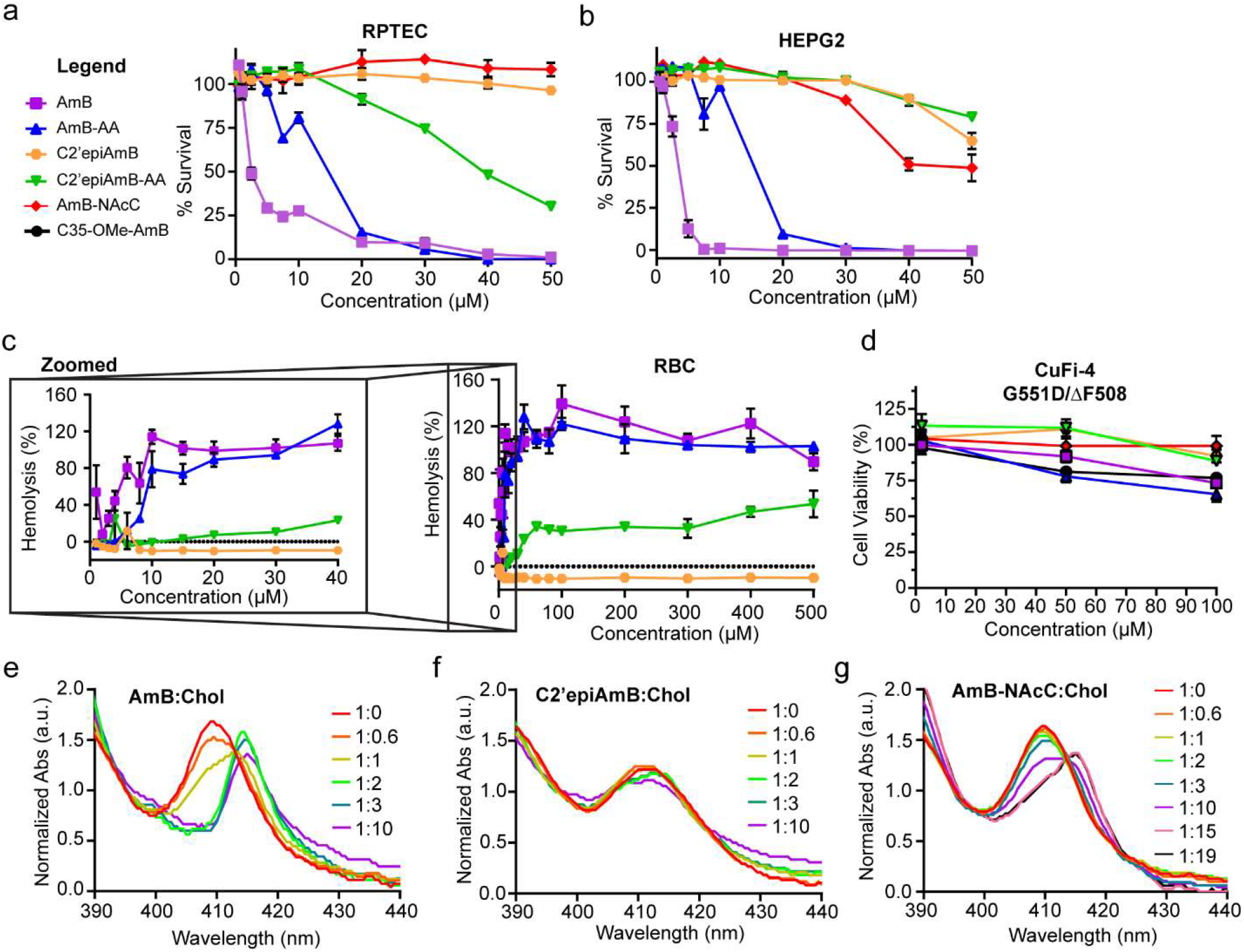
Toxicity profile of AmB-derivatives in cholesterol containing cell lines. **(a)** RPTEC (primary renal proximal tubule epithelial cells; n=3 biological replicates/conc.)) toxicity plot of AmB, AmB-AA, C2’epiAmB, C2’epiAmB-AA, AmB-NAcC. **(b)** HEPG2 (human liver cells; n=3 biological replicates/conc.) toxicity plot of AmB, AmB- AA, C2’epiAmB, C2’epiAmB-AA, and AmB-NAcC. **(c)** Hemolysis Plot of AmB, AmB-AA, C2’epiAmB, and C2’epiAmB-AA. **(d)** CuFi-4 Toxicity of AmB, AmB-AA, C2’epiAmB, C2’epiAmB-AA, AmB-NAcC, and C35-OMeAmB at 2, 50, and 100µM. **(e)** AmB:Cholesterol UV-Vis binding plot. **(f)** C2’epiAmB:Cholesterol UV-Vis binding plot. **(g)** AmB-NAcC:Cholesterol UV-Vis binding plot.

**Extended Figure 3:**
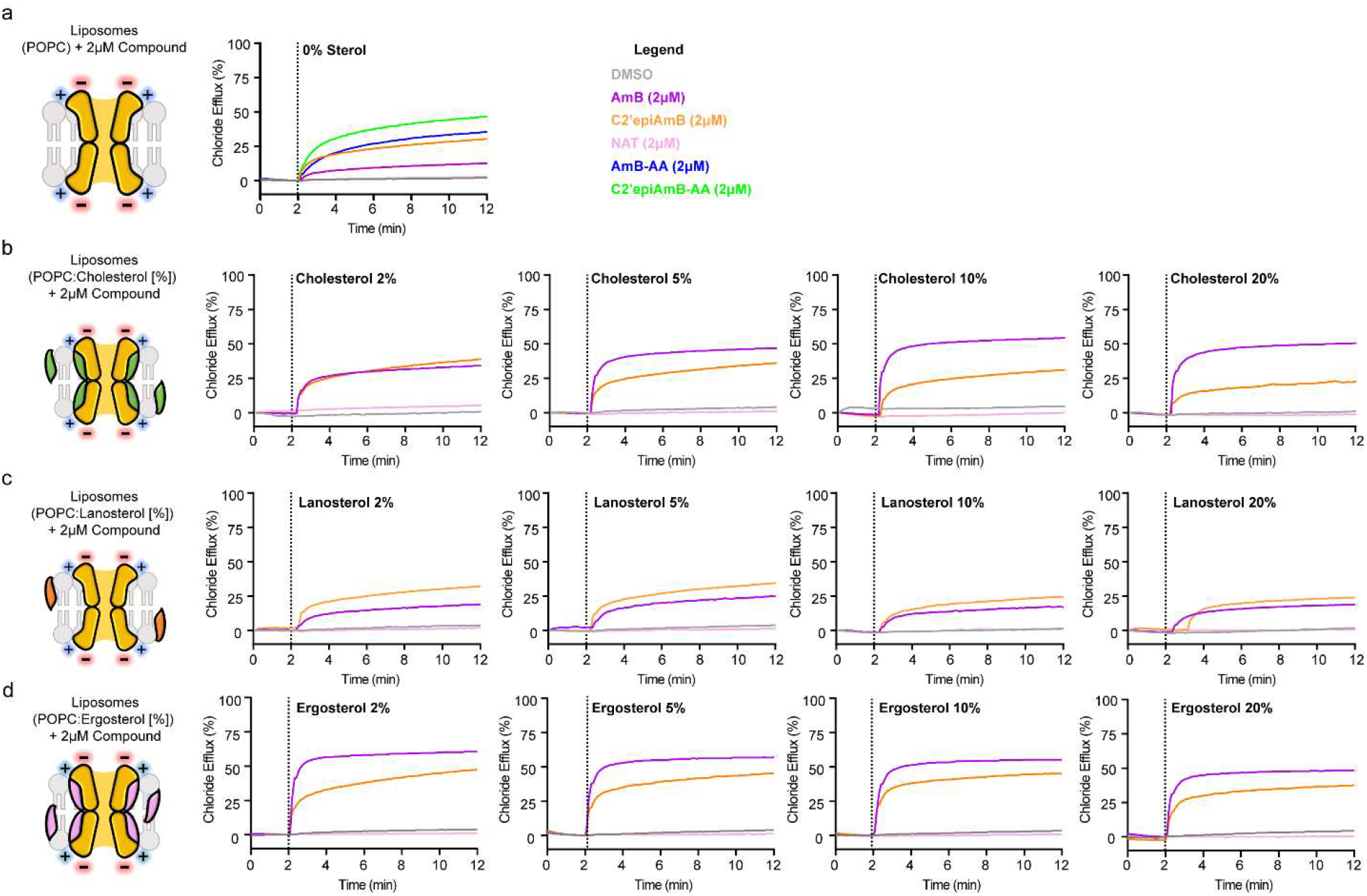
POPC Liposomes shows AmB channel formation in non-sterol containing membranes. **(a)** Chloride efflux of DMSO, AmB, C2’epiAmB, Natamycin, AmB-AA, and C2’epiAmB-AA (2 µM) in sterol free liposomes **(b)** DMSO, AmB, C2’epiAmB, and Natamycin (2 µM) in cholesterol containing liposomes (2, 5, 10, and 20%) **(c)** Chloride efflux of DMSO, AmB, C2’epiAmB, and Natamycin (2 µM) in lanosterol containing liposomes (2, 5, 10, and 20%) **(d)** Chloride efflux of DMSO, AmB, C2’epiAmB, and Natamycin (2 µM) in ergosterol containing liposomes (2, 5, 10, and 20%).

