## Supplemental Information for "Molecular prosthetics for CFTR designed for anion selectivity outperform amphotericin B in cultured cystic fibrosis airway epithelia"

**Reactions:** All manipulations of Amphotericin B were carried out under low light and stored under an anaerobic atmosphere at -20 °C. All reactions were monitored by RP-HPLC using an Agilent 1260 Infinity II series HPLC with 6530 LC/Q-TOF system equipped with an Ecplise XDB-C18 5 micron 4.6 x 150 mm column with UV detection at 406 nm.

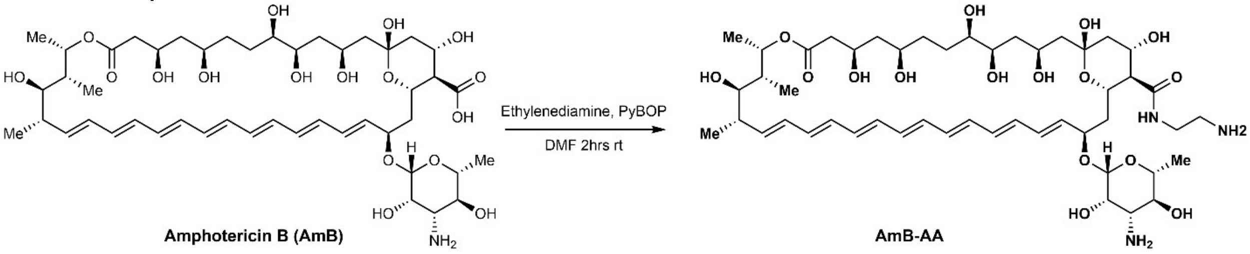


**Synthesis of AmB-AA**: The following protocol was followed and adapted from work by Maji, A. et.al[25]. Amphotericin B (250 mg, 0.27 mmol, 1 eq) was dissolved alongside ethylene diamine (54.2 µL, 0.81 mmol, 3 eq) in DMF (38 mM). To the mixture, $Et_3N$ (8.30 M) was added and stirred for 15 minutes at 25 °C. PyBop (134.8 mg, 0.405 mmol, 1.5 eq) was added to the solution and stirred for an additional 2 hours at 25 °C. The reaction was tracked via LC-MS for full conversion. Upon completion, the reaction mixture was slowly poured into stirring $Et_2O$ (250 mL per gram of amphotericin B used) and stirred for 10 minutes and then allowed to settle. The $Et_2O$ was partially decanted, and the precipitate was isolated via Buchner filtration using Whatman 50 filter paper and washed with $Et_2O$ to afford the desired product as an orange solid. Afterwards, the crude product was washed with EtOAc three times. For high purity material, the resulting material was dissolved in DMSO and purified via RP-HPLC (Waters SunFire Prep C18 OBD 5 micron 30 x 150 mm; 25 mL/min flow rate, MeCN: 2 mM $NH_4HCO_3$ in $H_2O$ 1:19 → 19:1 over 30 minutes). Material purity was then confirmed via LCMS at a wavelength of 406 nm to be >90% before biological testing (Eclipse XDB-C18 5 micron 4.6 x 150 mm; 0.5 mL/min flow rate, MeCN: 0.1% Formic acid in $H_2O$ 25:75 → 98:2 over 20 minutes).

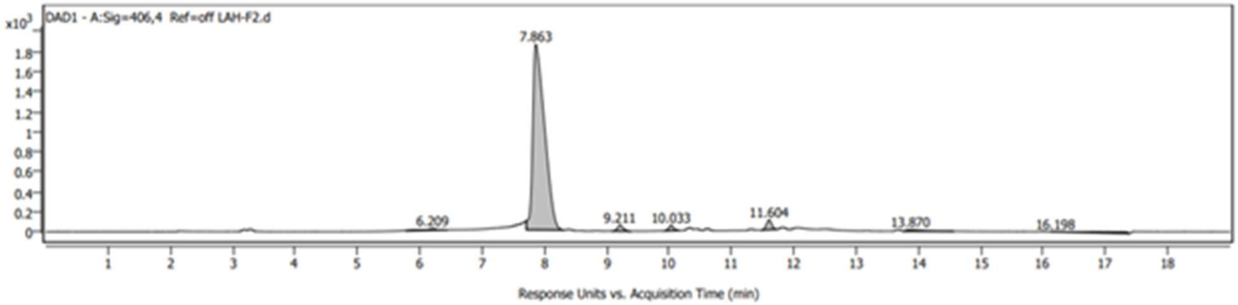


Calculated $C_{49}H_{79}N_3O_{16}$ $(M+H)^+$ : 966.5539
Found: 966.5669

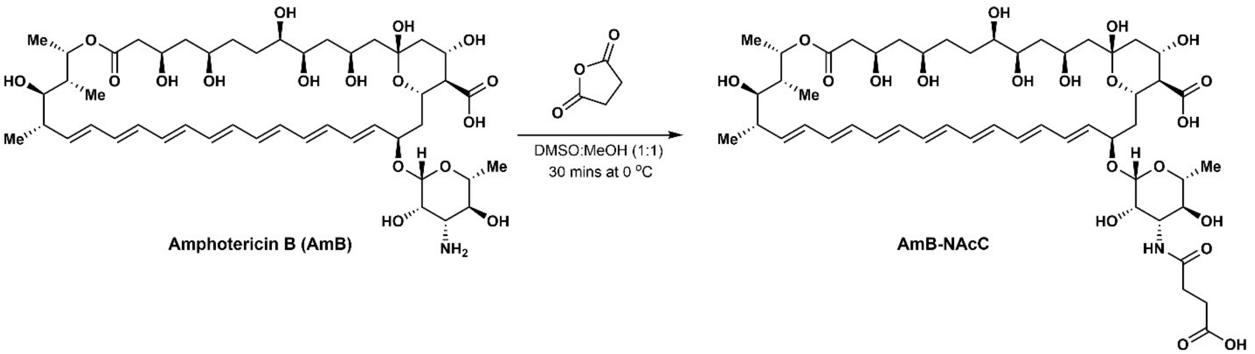


**Synthesis of AmB-NAcC:** The following protocol was performed and adapted from a previous report by Nicolaou, K.C. et.al[65]. Amphotercin B (1g, 1.08 mmol, 1eq) was dissolved in 10 mL of DMSO at room temperature and diluted with 10 mL of methanol. The reaction mixture was cooled in ice and succinic anhydride (95.1 µL, 1.17 mmol, 1.08 eq) was added dropwise while stirring. The reaction was set to stir as room temperature for 1 hour and tracked via LCMS for full conversion. Upon completion, the reaction mixture was slowly poured into stirring $Et_2O$ (250 mL per gram of amphotericin B used). The suspension was centrifuged for 5 minutes at 3700 rpm to isolate the solid orange-yellow product. The sample was dissolved in a minimal amount of DMSO and lyophilized overnight to obtain a dry, yellow powder. The yellow powder was purified by RP-MPLC (silica gel, 10 → 60% 10mM $NH_4HCO_3$ in acetonitrile). Material purity was then confirmed via LCMS at a wavelength of 406 nm to be >90% before biological testing (Eclipse XDB-C18 5 micron 4.6 x 150 mm; 0.5 mL/min flow rate, MeCN: 0.1% Formic acid in $H_2O$ 25:75 → 98:2 over 20 minutes).

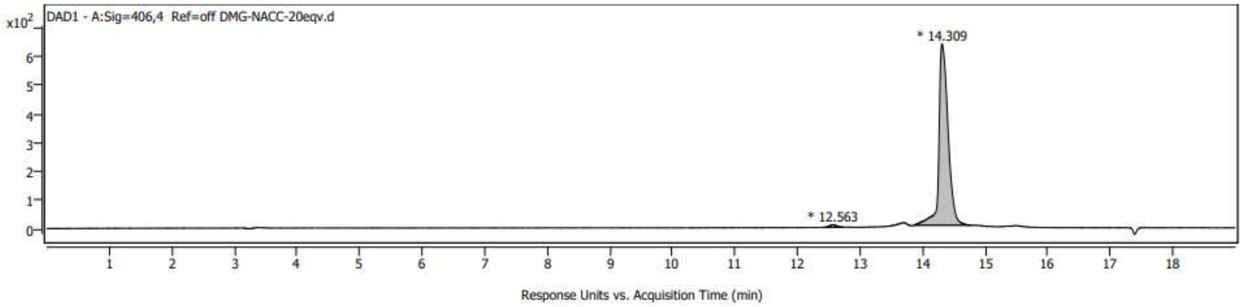


Calculated $C_{51}H_{77}NO_{20}Na$ $(M+Na)^+$ : 1046.4937
Found: 1046.4873

$^1$H NMR (500 MHz, PyrMeOD) δ 8.38 (s, 9H), 7.53 (t, *J* = 1.3 Hz, 9H), 7.13 (s, 15H), 6.41 (ddd, *J* = 17.0, 14.7, 10.6 Hz, 2H), 6.20 (dtd, *J* = 11.6, 8.7, 4.6 Hz, 6H), 6.17 – 6.03 (m, 4H), 5.32 – 5.21 (m, 2H), 5.19 (s, 49H), 4.67 (d, *J* = 13.0 Hz, 2H), 4.63 – 4.51 (m, 2H), 3.22 (p, *J* = 1.6 Hz, 10H), 2.58 (q, *J* = 6.5 Hz, 4H), 2.40 – 2.23 (m, 4H), 2.18 – 2.08 (m, 2H), 1.83 – 1.77 (m, 2H), 1.77 – 1.71 (m, 2H), 1.70 – 1.55 (m, 3H), 1.52 – 1.26 (m, 7H), 1.22 (d, *J* = 6.1 Hz, 3H), 1.14 (d, *J* = 6.4 Hz, 3H), 1.03 (d, *J* = 6.4 Hz, 3H), 0.96 (d, *J* = 7.1 Hz, 3H).

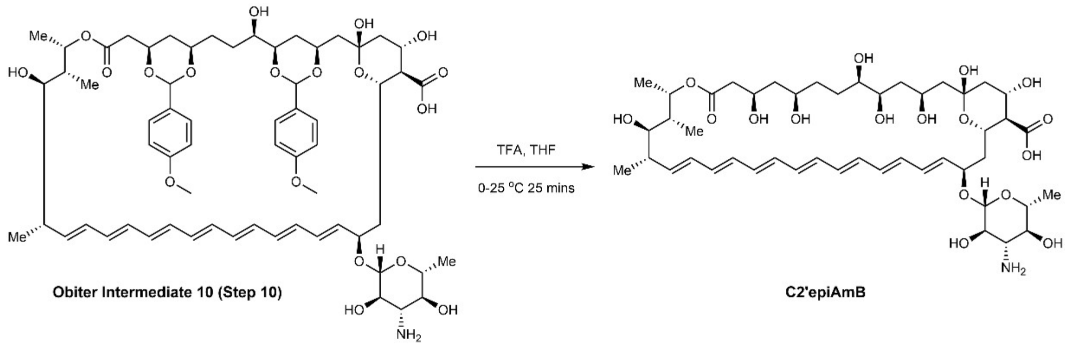


**Synthesis of C2’epiAmB**: This material was synthesized using a precursor provided by Obiter (Obiter Intermediate 10; **Step 10**) following the reported procedure in Maji, A. et.al[25]. To a reaction flask 7.10 mL (60 mM) of THF and 4.73 mL (90 mM) of Water were mixed (3:2) and cooled to 0 °C. To this stirring solution, 1.18 mL (13.3 µmol, 31.30 eq) of trifluoromethanesulfonic acid was slowly added and allowed to cool back to 0 °C. Once cooled, 500 mg (425.75 µmol, 1 eq) of Step 10 material was added. The reaction mixture was sonicated until solid material was fully dissolved and allowed to cool back to room temperature. The reaction was run at room temperature for 23 minutes then cooled back to 0 °C for 2 minutes. Once cooled 4.11 mL (29.5 mmol, 69.3 eq) of $Et_3N$ was added. The reaction mixture was diluted in a mixture of diethyl ether and acetone (~1:1). If material does not crash out, and two layers were still visible, then slightly more acetone was used until a cloud of precipitate was seen. The crashed-out material was then centrifuged at 3800 rcf for 5 minutes. The supernatant was decanted, and the solid material was washed again with diethyl ether and acetone mixture (1:1) and the process was repeated until minimal amount of yellow supernatant was seen. This process afforded a yellow solid C2’epiAmB (average yield was >88%). Material could be used or carried forward without further purification. Compound purity (>90%) and reaction completion was determined via LCMS using an Eclipse XDB-C18 5 micron 4.6 x 150 mm column, using a gradient of Water:ACN (+0.1% formic acid) starting at 95:5 for 1 minute and increased to 5:95 over a period of 10 minutes followed by a return to 95:5 over a period of 2 minutes.

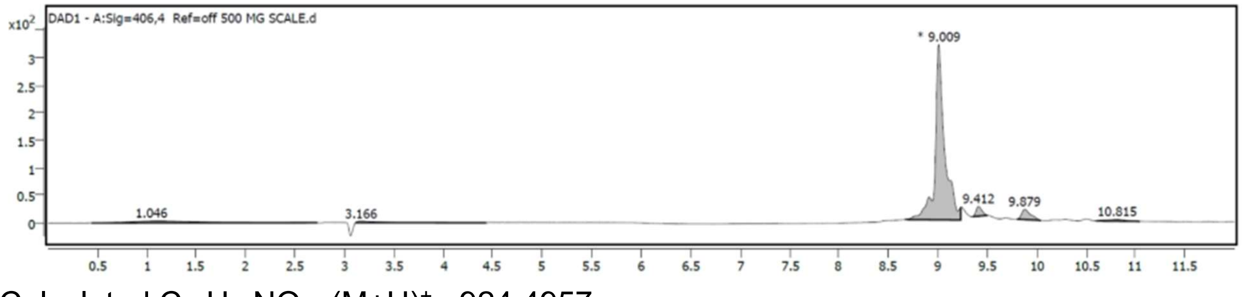


Calculated $C_{47}H_{73}NO_{17}$ $(M+H)^+$ : 924.4957
Found: 924.5056

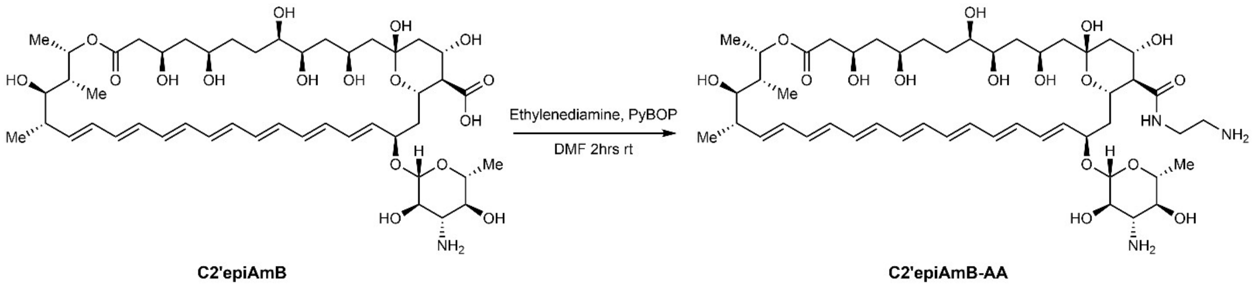


**Synthesis of C2'epiAmB-AA:** The following protocol was followed and adapted from previous reports[36]: Dissolve 100 mg (103.89 µmol, 1 eq) of **C2'epiAmB** in 519 µL of DMF (200 mM). To the mixture add 296 µL (350 mM) of $Et_3N$ and stir for 5 minutes at 25 $^oC$. To the mixture 270.31 mg (519.43 µmol, 5 eq) of PyBop was added to the solution and stirred for an additional 5 minutes at 25 $^oC$. The solution was also sonicated until all materials were fully dissolved. To this reaction mixture 41.67 µL (623.32 µmol, 6 eq) of diethyleneamine was added and the reaction mixture was allowed to stir at room temperature for 30 minutes. Upon completion, the reaction mixture was slowly poured into stirring $Et_2O$ (25 mL), the mixture was centrifuged at 3800 RCF for 5 minutes at 0 $^oC$. The supernatant was decanted and to the remaining black/orange solid 25 mL of EtOAc was added and mixture was sonicated before being centrifuged at 3800 RCF for 5 minutes at 0 $^oC$. The supernatant was decanted and solid dried under vacuum to afford the desired product as a yellow/orange solid. For pure material (>90%), the resulting material was dissolved in DMSO and purified via RP-HPLC (Waters SunFire Prep C18 OBD 5 micron 30 x 150 mm; 25 mL/min flow rate, MeCN: 2 mM $NH_4HCO_3$ in $H_2O$ 1:19 → 19:1 over 30 minutes). Material purity was then confirmed via LCMS at a wavelength of 406nm following the protocol shown for **C2'epiAmB**.

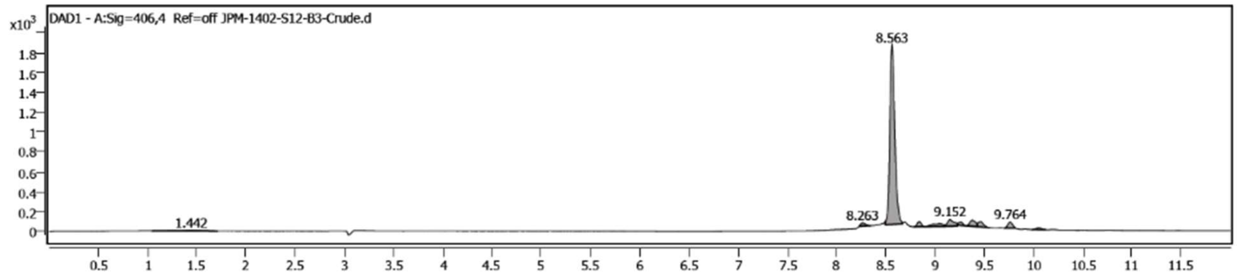


Calculated for $C_{49}H_{79}N_3O_{16}$ $(M+H)^+$: 966.5539
Found: 966.5713
